# Harnessing olfactory bulb oscillations to perform fully brain-based sleep scoring and track sleep/wake transitions across multiple time scales in mice

**DOI:** 10.1101/109033

**Authors:** Sophie Bagur, Marie Masako Lacroix, Gaëtan de Lavilléon, Julie M Lefort, Hélène Geoffroy, Karim Benchenane

## Abstract

It has long been thought that sleep scoring could not be achieved with brain signals alone despite the deep neuromodulatory transformations that accompany sleep state changes. Here we demonstrate using multi-site electrophysiological LFP recordings in freely moving mice that gamma power in the olfactory bulb (OB) allows for clear classification of sleep and wake. Coupled with hippocampal theta activity, it allows the construction of a sleep scoring algorithm that relies on brain activity alone. This method reaches over 90% homology with classical methods based on muscular activity (EMG) and video tracking. Moreover, contrary to EMG, OB gamma power allows correct discrimination between sleep and immobility in ambiguous situations such as fear-related freezing. We use the instantaneous power of hippocampal theta oscillation and OB gamma oscillation to construct a 2D phase-space that is highly robust across mice and days. Dynamic analysis of trajectories within this space yields a novel characterization of sleep/wake and wake/sleep transitions as deeply divergent phenomena. Whereas waking up is a fast and direct transition, falling asleep is best described as stochastic and gradual change. Altogether this methodology opens the avenue for multi-timescale characterization of sleep states with high temporal resolution based on brain signals only.

## Introduction

A current challenge in neuroscience is defining the different states of brain activity and describing how they impact the computations performed by neural networks. The most dramatic change of state is that between sleep and wakefulness that involves modifications of cortical activation (Steriade *et al.*, 1993), gene expression (Cirelli & Tononi, 2000), engagement with the outside world and clearance mechanisms (Xie *et al.*, 2013) amongst other changes throughout the whole organism (Benington & Heller, 1995; Imeri & Opp, 2009). Despite these profound transformations, to date, we surprisingly lack an easily measured marker of brain activity that allows unambiguous, moment-to-moment identification of sleep and wake states.

Several sleep scoring methods have been proposed in human research (Rechtschaffen & Kales, 1968; Iber *et al.*, 2007) and are now widely accepted by the scientific community. There is on the contrary no clear consensus on sleep scoring procedures in the rodent (Datta & Hobson, 2000). The methods used are generally completely manual or rely on manually scored training data to calibrate automatic algorithms (Table 1), therefore adding to the variability between laboratories the problem of inter-scorer variability.

Moreover all current sleep scoring methods essentially rely on motor activity to discriminate sleep from wake, see Table 1 (Veasey *et al.*, 2000; Louis *et al.*, 2004; Crisler *et al.*, 2008; Gross *et al.*, 2009; Stephenson *et al.*, 2009; Brankack *et al.*, 2010; Rytkönen *et al.*, 2011; Liang *et al.*, 2012; Zeng *et al.*, 2012). This renders all methods inherently vulnerable to any mismatch between these brain states and the level of motor activity such as during freezing, a commonly-used behaviour in mice or any sleep anomalies causing movement during sleep (Schenck & Mahowald, 2002).

The state of the art therefore presents both a conceptual and a technical problem regarding the definition of sleep and wake.

Sleep scoring methods are based on the assumption that the information about sleep states is contained in the recorded signal and can be used as a marker (Libourel *et al.*, 2015). In order to implement a reliable sleep scoring, the candidate marker of sleep and wake must not only show a strong average difference between the two states but this change must be systematic and sustained throughout each state, with a clear separation between the values in each state. A bimodal distribution with good separation of the two component distributions is the optimal situation to allow moment-by-moment discrimination.

Despite multiple studies, such a clear-cut situation has never been found when using brain signals (Brankack *et al.*, 2010). Therefore attempts to identify sleep with brain signals only rely on more elaborate methods that extract composite features from LFP data (Gervasoni *et al.*, 2004). The first step of dimensionality reduction is highly dependent on the data at hand and often results in mapping data onto axes which are highly variable between animals and sessions, requiring post-hoc human labelling procedures and providing low separation between states (Gervasoni *et al.*, 2004). To compensate for the poor quality of the brain-related sleep markers, machine learning techniques have been used in several studies but with no decisive improvement (Crisler *et al.*, 2008; Yu *et al.*, 2009; Rytkönen *et al.*, 2011; Chou *et al.*, 2013).

Here we propose a novel brain-related marker allowing to reliably track transitions from sleep to wakefulness. Indeed the gamma power (50-70Hz) measured in the olfactory bulb has been shown to vary between sleep states (Manabe & Mori, 2013) but we show here that it displays the desired characteristics to continuously identify the different sleep states. We thus propose that these oscillations can be used as a direct read-out of the brain network responsible for sleep/wakefulness cycles on the time scales both of stable brain states and fast state transitions.

Accordingly, we show that the gamma oscillation in the olfactory bulb is strongly suppressed during sleep and continuously present during waking. Moreover, the distribution of the gamma power follows a bimodal distribution that is the optimal situation for an automatic separation procedure. Coupling this indicator with the classical hippocampal theta/delta power ratio allows us to construct a fully automated sleep scoring algorithm that classifies wake, rapid eye movement sleep (REM) and non-REM sleep (NREM) based on brain state alone.

We then use these variables to construct a robust 2D phase-space that is highly robust across mice and days. This phase space forms the basis of an innovative analytical methodology for the study of fast time scale transitions using kinematic modelling of trajectories. As an application, we provide strong evidence for a deep asymmetry between transitions from sleep to wake and wake to sleep. In particular, the awakening transition displays strongly driven, fast dynamics whereas the process of falling asleep is more stochastic and slower.

Together, we propose a new sleep scoring method that has strong methodological (automatic, reproducible, no training data required) and conceptual (no reliance on motor activity) advantages over traditional methods and allows us to characterize the fine dynamics of state transitions in the brain.

## Results

### Olfactory bulb gamma power modulation throughout brain states

Classical sleep scoring methods differentiate sleep and wake states using EMG activity or the animal’s motion recorded using accelerometers or video tracking (see Table 1, Veasey *et al.*, 2000; Louis *et al.*, 2004; Crisler *et al.*, 2008; Gross *et al.*, 2009; Stephenson *et al.*, 2009; Brankack *et al.*, 2010; Rytkönen *et al.*, 2011; Liang *et al.*, 2012; Zeng *et al.*, 2012). Theta and delta power recorded in the hippocampus or cortex (due to the volume conduction of theta oscillations) can then be used to discriminate REM from NREM sleep. According to classical sleep scoring methods, wake is defined by high EMG activity and low HPC signal during arousals, irregular HPC activity during quiet wake or theta oscillation during active exploration. On the contrary, sleep is defined by low EMG power and NREM is discriminated from REM using the theta/delta power ratio in the hippocampus. During NREM delta power is strong whereas highly regular theta oscillations are observed during REM (Fig. 1A).

**Figure 1.**
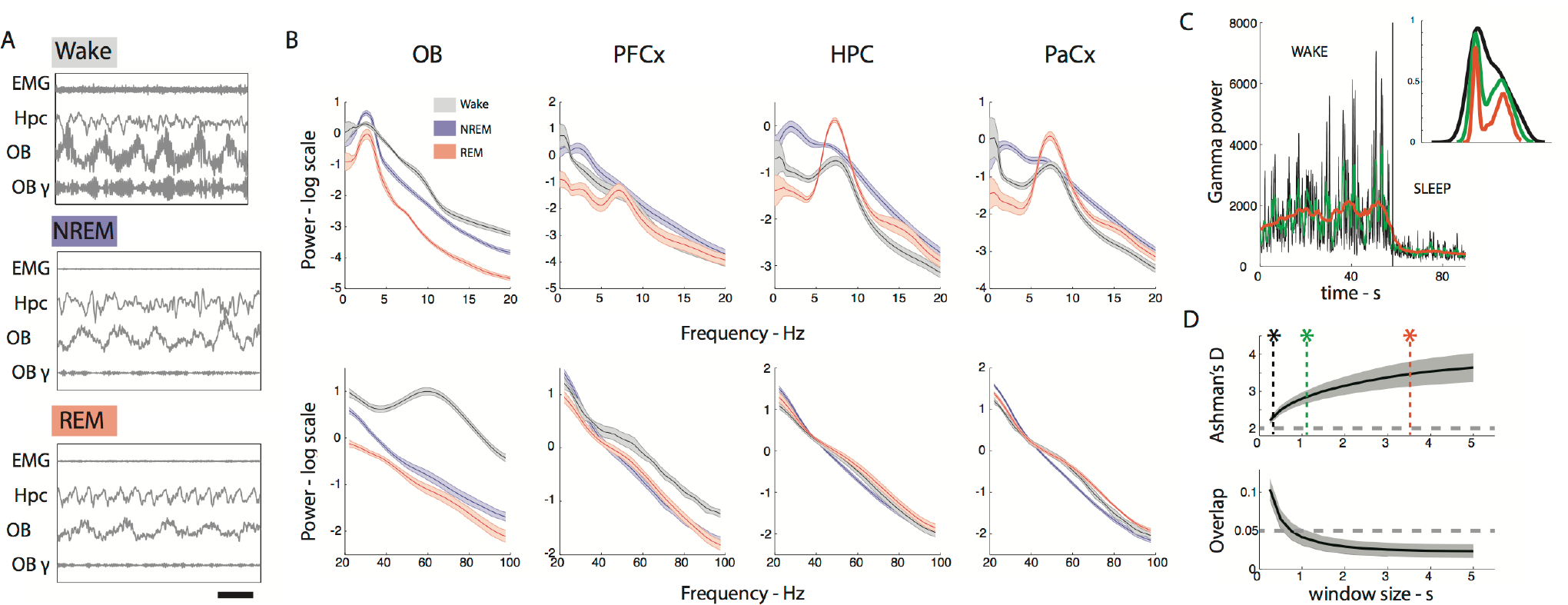
**A**. Example data showing EMG, HPC and OB activity in three brain states. Wake is characterized by high EMG activity and high gamma power in the OB whereas sleep is characterized by low EMG activity and low gamma power in the OB. HPC LFP shows regular theta activity during REM sleep. Filtered signals from the OB in the gamma band (OB-g, 50-70Hz) shows the remarkable decrease in gamma during sleep states. (Scale bar: 0.5s) **B**. Low (top) and high (bottom) frequency spectra from different brain regions during NREM, REM and wake states as classified using movement based scoring (EMG or filmed activity). Note the strong increase in gamma activity in the OB in the wake state. (n=10 for OB and HPC, n=6 for PFCx and PaCx, error bars: s.e.m) **C**. Gamma power in OB is plotted as a function of time as the animal transitions from wake to sleep and the distribution of the corresponding values is shown on the right. The data has been smoothed different window lengths (0.1, 1 and 4s respectively). The fast fluctuations present in the awake state are smoothed out as the window size increases yielding a more clearly bimodal distribution with larger smoothing window. **D**. Ashman’s D (bimodality indicator, significant if larger than 2) (i) increases and the overlap between the two gaussians (ii) decreases with the length of the smoothing window. The stars show window sizes illustrated in C.

**Table 1.**
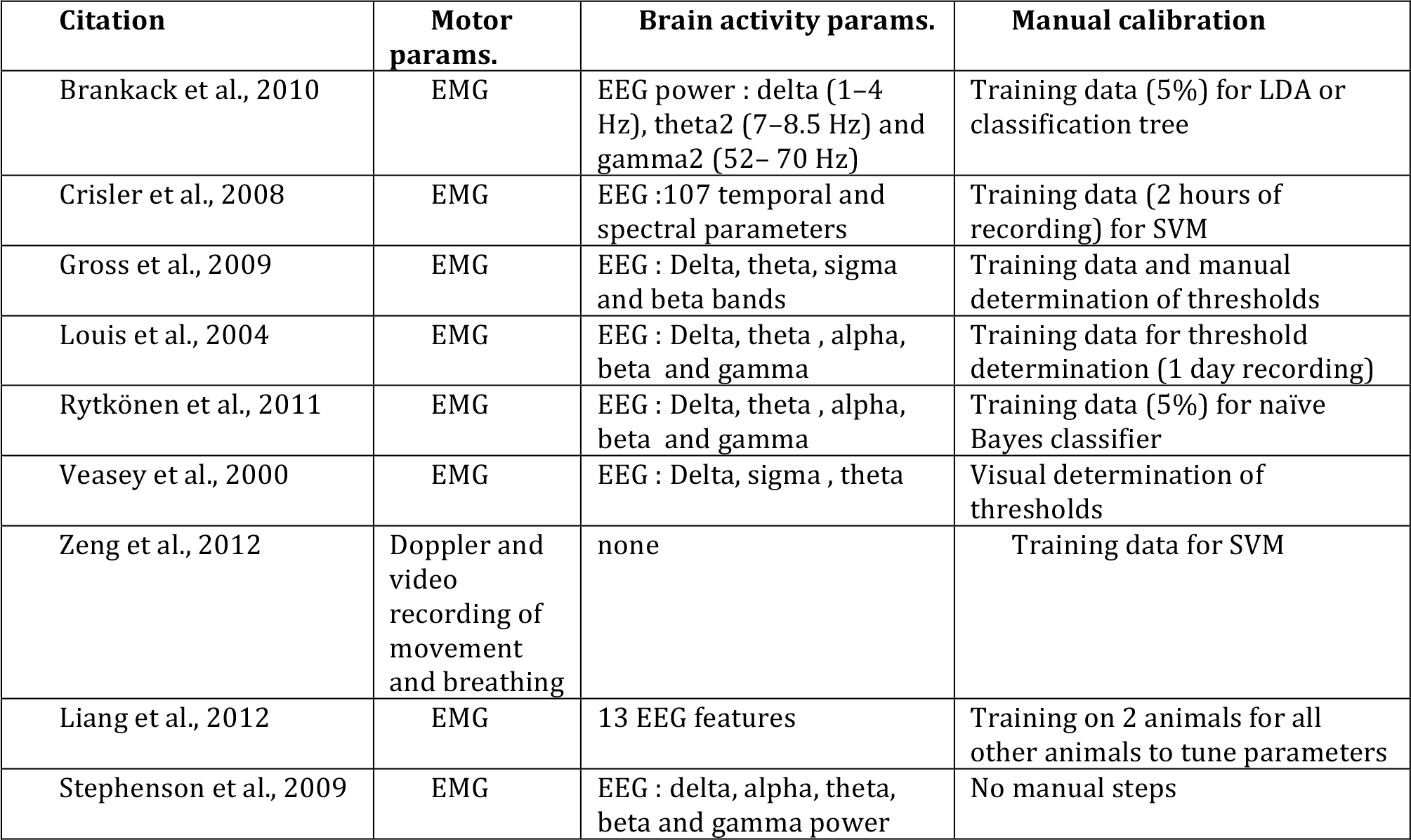

In order to construct a sleep scoring method that relies on brain signals alone, we screened multiple brain regions to find a good predictor for discriminating between sleep and waking states. We recorded from multiple brain regions in 15 freely-moving mice: the olfactory bulb (OB, n=15), the hippocampus (HPC, n=15), the prefrontal cortex (PFCx, n=6) and the parietal cortex (PaCx, n=6) cortex. Mice were recorded for an average of 6.6 ± 0.58 hrs (minimal recording length: 2hrs) in their homecages in the light period and slept on average 58% of the time. We initially used a classical sleep scoring method based on movement and hippocampal activity to establish a database of recordings from different brain states using 10 of the 15 mice that either were implanted with an EMG wire in the nuchal muscles (n=6) or tracked using video (n=4).

The average spectra over wake, NREM and REM periods are shown in Fig. 1B. In cortical and hippocampal areas, as expected, REM and NREM showed strong differences in the theta and delta band and wake periods showed less low frequency power. In cortical and hippocampal areas, no individual frequency band allows to discriminate well between sleep and wake.

Remarkably however, we found a strong increase in power in the OB during waking relative to sleep states. This difference was strongest in the low gamma band centred at 60Hz as previously described (Manabe & Mori, 2013). This change is clear in the Fig. 1A which shows OB activity that is constantly modulated by the breathing cycle but which displays a sustained faster oscillation only in the wake state. Crucially, gamma power was low in both sleep states, suggesting that this parameter could replace muscle activity for discriminating wake from REM sleep. We also observed that beta activity in the OB was a good predictor of REM vs NREM sleep, reinforcing this striking relationship between OB oscillations and brain states (Fig S1) but focussed on the gamma activity as a candidate replacement for muscular activity in sleep scoring.

OB gamma power displays strong fluctuations correlated with breathing activity on the scale of around a second (Manabe & Mori, 2013). To find the appropriate time scale for tracking the changes in gamma power related to brain state changes, we applied a smoothing window of varying length to the instantaneous gamma power. As the smoothing window increased in length, the distribution of gamma power became more distinctly bimodal and the two underlying distributions clearly separated (Fig. 1C-D). We found that smoothing windows larger than 1s produced two normal distributions overlapped by less than 5% (Fig. 1D bottom). This analysis allowed us to establish a set of parameters (frequency, smoothing window) that establish gamma power in the OB as a promising predictor for discriminating between wake and sleep on fine timescales of the order of one second without any reliance on muscular activity.

### Construction and validation of the sleep scoring algorithm

A schematic of the sleep scoring algorithm is shown in Fig. 2A. All steps are automatic and do not require any supervision by the user. Instantaneous smoothed gamma power in the OB shows a bimodal distribution that can be well fit by a sum of two Gaussian functions (Fig. 2B) (mean R^2^ = 0.98 ± 0.009). The two component distributions correspond to gamma power during sleep and wake periods defined by movement. Since the amplitude of these distributions depends on the proportion of time spent in each state, they are normalized (see Methods) and the sleep/wake threshold is defined as the intersection of the two Gaussian curves (Fig. 2Bi). Below threshold values of gamma power are defined as sleep and above threshold values as wake.

**Figure 2.**
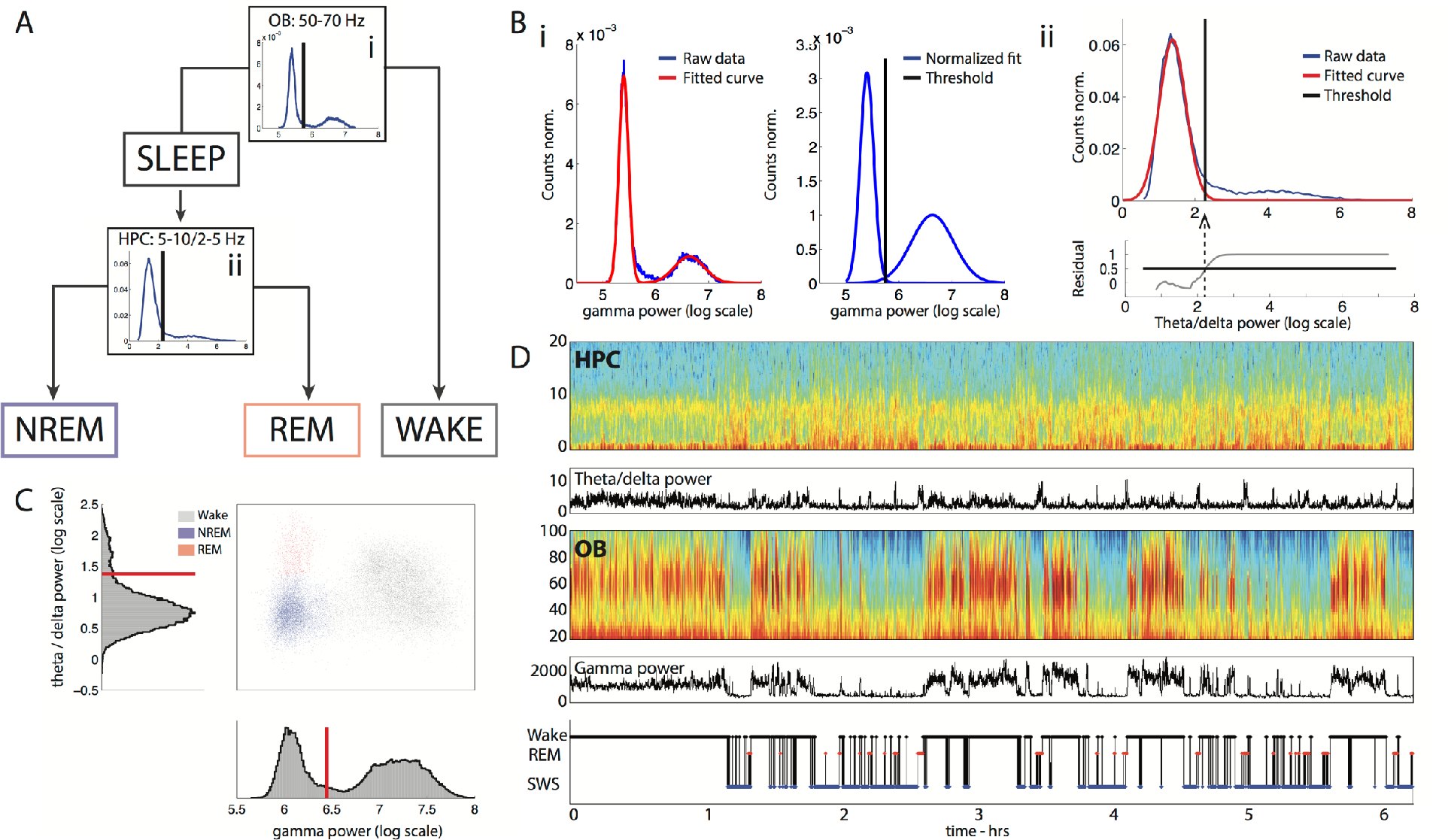
**A**. *Flowchart of data through the scoring algorithm. Sleep and wake states are first classified based on OB gamma (i). Sleep data is then further classified into REM and NREM based on HPC theta / delta power ratio (ii)*. **B**. *Example of automatic thresholding of distributions. (i) Two gaussian distributions are fit to the distribution of OB gamma power (left) and their areas are equalized (right). The threshold is placed at the intersection of the two distributions. (ii) A gaussian distribution is fitted to the distribution of HPC theta / delta power ratio during sleep. The residuals are shown in the bottom plot. The threshold is placed at the point where the fit explains less than 50% of the data*. **C**. *Example 2D phase space. Each 3s period of recording is plotted according to its average OB gamma power value and average HPC theta / delta power ratio showing the three brain states identified: NREM (blue), REM (red) and Wake (grey). Corresponding histograms are shown along the relevant axis with automatically determined thresholds in red*. **D**. *Example data set showing HPC low frequency spectrogram (i) with theta / delta power ratio below and OB high frequency spectrogram (iii) with gamma power below. Hypnogram is shown at the bottom*.

Each time point is now attributed to one of the three states, based on its OB gamma power and HPC theta/delta ratio (Fig. 2C). Brief periods of less than 3s are merged with the neighbour states (see methods for details). An example session is shown in Fig. 2C-D that illustrates the construction of a two dimensional phase space for brain states (Fig. 2C). This space demonstrates the clear separation of brain states even after the merging and dropping of short epochs. This shows that the continuity hypothesis does not lead to any aberrant classification.

We validated the sleep scoring algorithm by comparing it to manual sleep scoring performed using HPC LFPs and EMG activity, the classical golden standard. Two expert scorers independently scored sessions from 4 mice with an average inter-scorer overlap of 89 ± 3% and Cohen’s K of 0.81. On average the automatic and manual sleep scoring overlapped by 90 ± 2% (Cohen’s K: 0.83) throughout the sessions (Fig. 3A).

**Figure 3.**
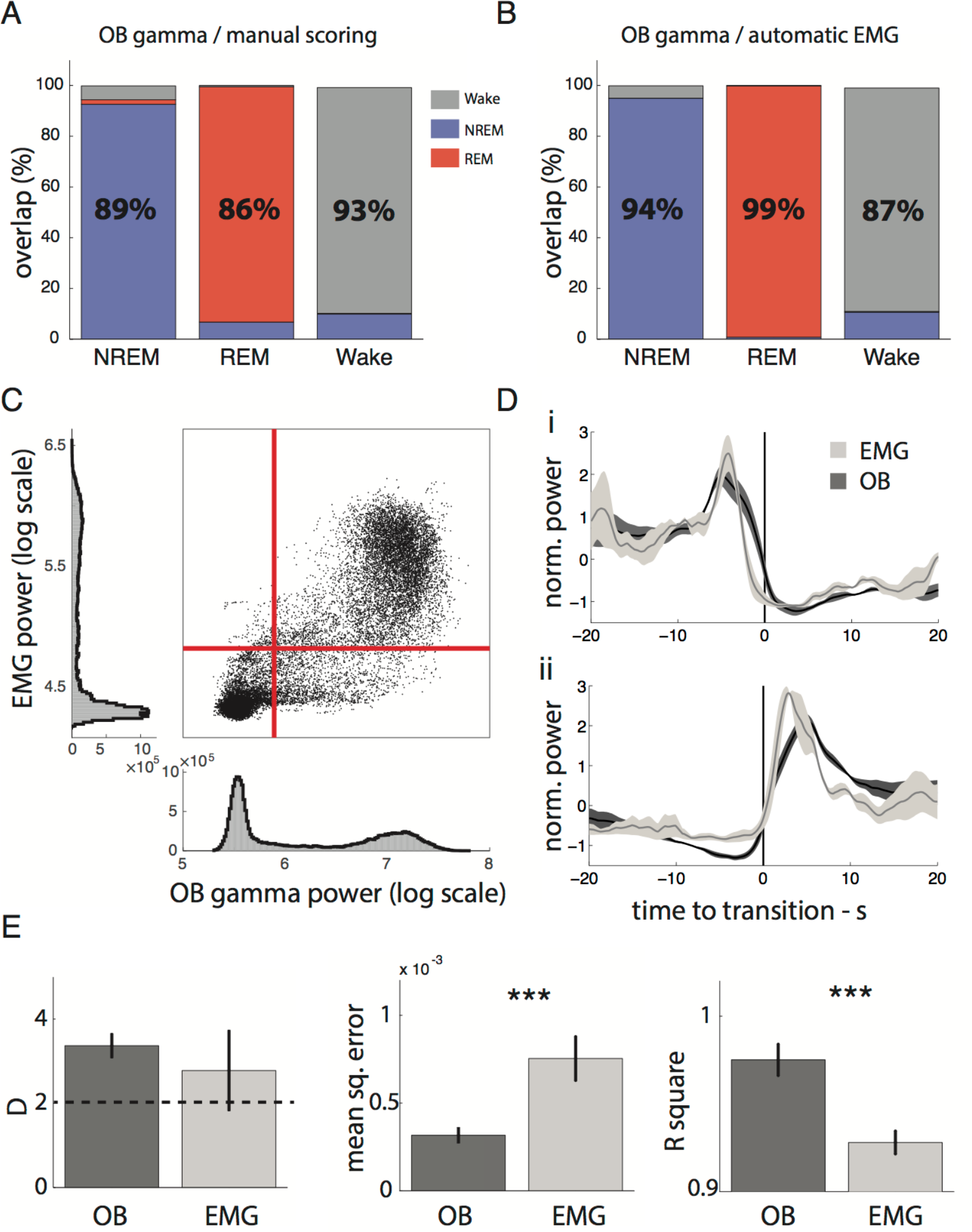
**A**. *Overlap of manual scoring with OB gamma scoring. Each column gives the percent of manually identified brain state (x label) that is classified as NREM (blue), REM (red) and wake (grey) respectively using OB gamma scoring. (n=6)* **B**. *Overlap of automatic EMG scoring with OB gamma scoring. (n=6)* **C**. *Correlation of OB gamma power and EMG power for an example mouse with automatically determined thresholds for each distribution in red. Dots in the upper right and lower left hand corner are identically classified by both approaches*. **D**. *OB gamma and EMG power triggered on transitions from wake to sleep and sleep to wake determined by the OB gamma demonstrating tight temporal locking of changes with both indicators (n=6)*. **E**. *Comparison of the fit quality of bimodal distribution of two gaussians to gamma power and EMG power distributions. Both distributions are strongly bimodal as the average Ashman’s D is larger than 2. However both mean square error and Rsquare show that gamma power distributions are better fit by a sum of gaussians. (n=15 for gamma, n=6 for EMG. Ttest,=p=0.43, 0.009,0.0005 respectively)*

To more systematically compare the two approaches used to distinguish wake from sleep, gamma power in the OB and EMG power, we also performed scoring using an automatic EMG scoring algorithm (see Methods). Agreement between the two approaches was 93% (Cohen’s K: 0.85) (Fig. 3B) on average and the two signals were highly correlated at all times and time-locked at transition points (Fig. 3C,D).

We compared how well the distributions of each variable were described by fitting with two Gaussians (Fig. 3E). Both variables were strongly bimodal (Fig. 3E, *left*), however the error of the fit is higher for the EMG power. We found that this error was explained by a higher proportion of values in the trough between the two Gaussians (11 ± 2% for EMG and 4 ± 3% for gamma power). This indicates that the ‘ambiguous’ zone between sleep and wake is more densely occupied when using EMG scoring leading to more potential errors.

This demonstrates that sleep scoring using gamma power in the OB and EMG, using either automatic or manual methods, give very similar classification of brain states, confirming that gamma power is a good predictor of wake and sleep as classically defined. Moreover, gamma power provides distributions with a clearer separation than EMG power, making it a more reliable predictor.

### Robustness of the sleep scoring method

Sleep scoring is often performed on large batches of animals, requiring simple surgeries and a high success rate. It is known that theta rhythm can be easily recorded in the hippocampal area using a single LFP wire. How does gamma power used for sleep scoring depend on the exact placement of the recording site? To answer this question, we simultaneously recorded activity from multiple depths in the olfactory bulb covering the outer and inner plexiform layers, the mitral cell layer and granular cell layer using a sixteen sites linear probe (Fig. 4A). We found that gamma oscillations could be observed at all depths and sleep scoring performed using electrodes at all depth highly overlapped (>92%) with classical, movement based sleep scoring (Fig. 4B). We however observed that the separation between wake and sleep peaks was best in the deeper recording sites and in particular the most coherent scoring was found in those sites within the granule cell layer where gamma oscillations are visibly stronger (Fig. 4C,D).

**Figure 4.**
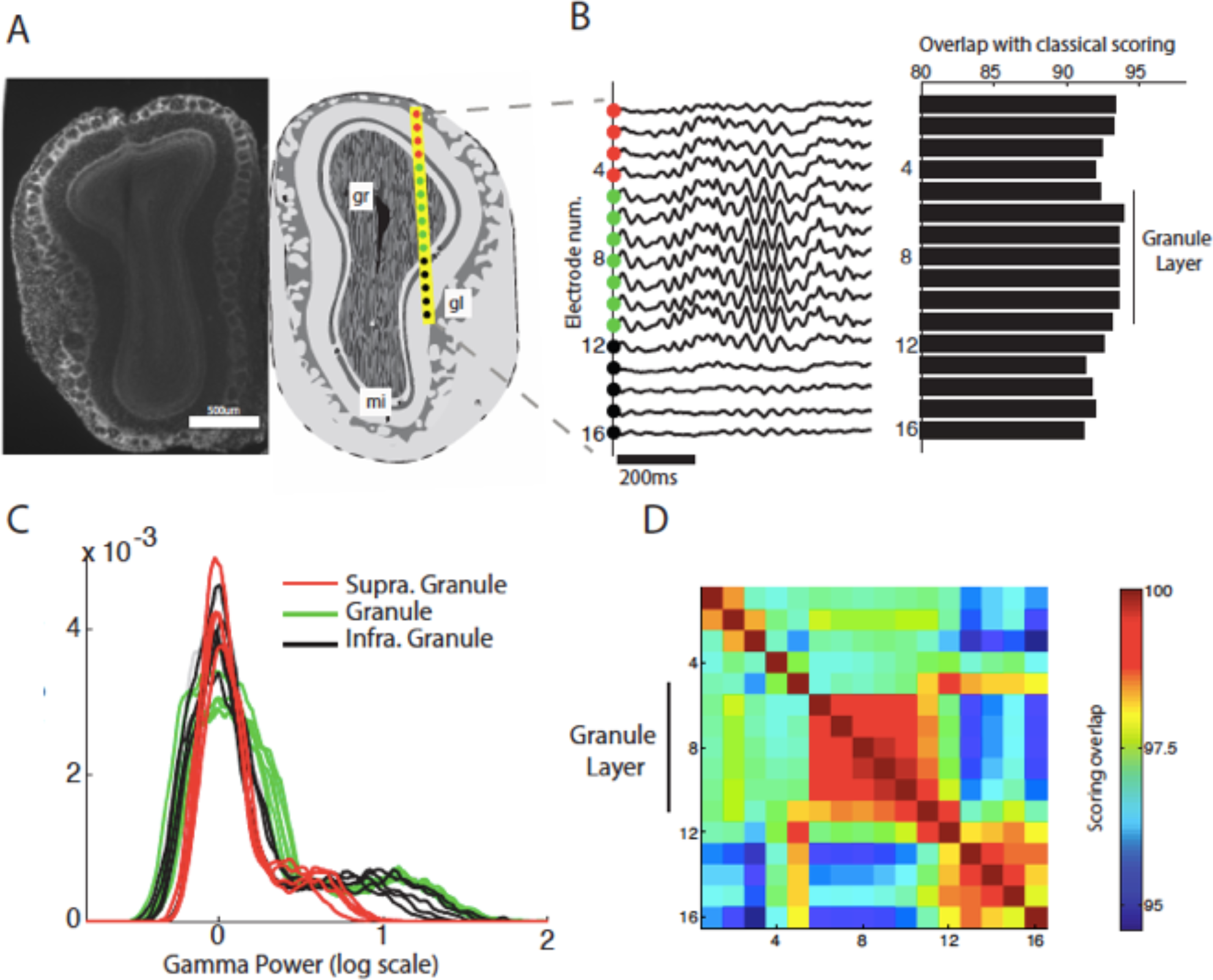
**A**. *Anatomical position (right) of 16-site silicon probe in the olfactory bulb as estimated from histological examination (left). This allows to estimate that sites 1-4 are above the granule layer and sites 12-16 are below*. **B**. *Sleep scoring is performed using the gamma activity from each electrode site and compared to sleep scoring using the animal’s movement. Accuracy is calculated as total overlap in sleep/wake periods*. **C**. *Gamma power distributions for each electrode separated into supragranular, granular andinfragranular layers. Note that the strong separation of sleep and wake peaks is clearest in electrodes within and below the granule layer*. **D**. *Correlation matrix of sleep scoring performed using gamma power from different depths. Each square shows the percent overlap between scoring performed with the corresponding electrodes. All values arehigh (above 95%) but the granule cell layer shows particularly coherent scoring (>99%)*.

This demonstrates that placement of the LFP wire for reliable scoring does not require great precision during implantation, assuring good scoring for all implanted animals. The granule cell layer however appears to be the optimal anatomical region to ensure reliable scoring since it shows the highest coherence in gamma power fluctuations. The coordinates we recommend aim for the center of this zone (AP +4, ML +0.5, DV −1.5).

An optimal sleep scoring technique must provide easily comparable results in the same animals throughout time and between animals. In other words, the phase space used to define sleep states must be stable. This phase space was constructed so that the separation between wake and sleep on the one hand and REM and NREM on the other hand used orthogonal axis. This simple space is remarkably consistent among animals and across days as can be seen by the similar position of the clouds of points representing each state (Fig. 5A).

**Figure 5.**
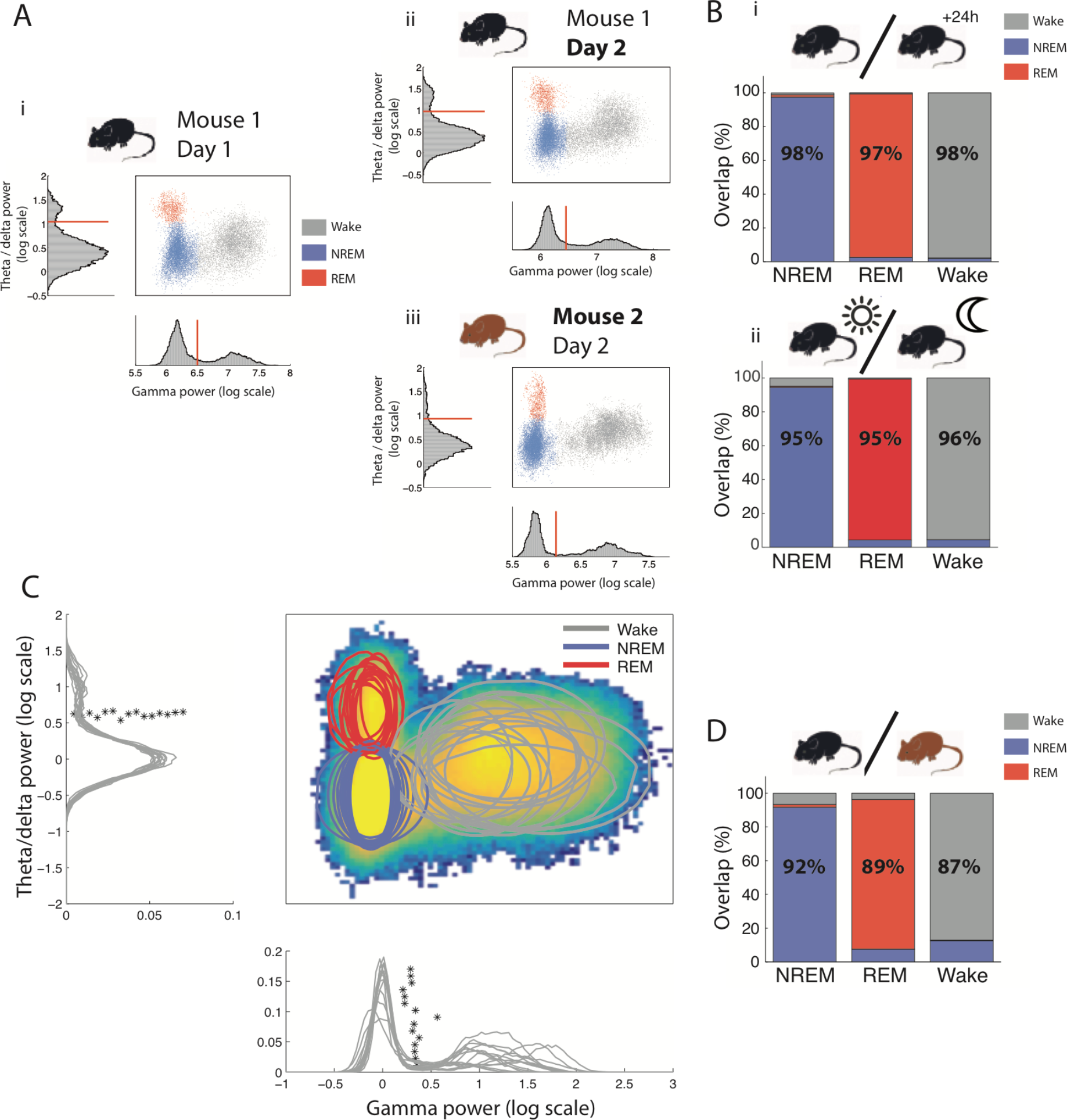
**A**. *Phase space of brain states and corresponding histograms along the relevant axis with automatically determined thresholds of the same mouse over different days (i vs ii) and in two different mice (i vs iii) demonstrating the highly conserved architecture across time and individuals*. **B**. *Overlap of scoring using thresholds determined on the same mouse on one day and applied the following day (i) and during successive light and dark periods (ii). Each column gives the percent of the brain state (x label) identified using thresholds determined on the reference data set that is classified as NREM (blue), REM (red) and wake (grey) using thresholds determined using data from different days (i) or during the light period (ii). (n=10mice were recorded on consecutive days and used for inter-day scoring, n=5 mice were recorded over a 24h period of light and dark periods)* **C**. *Heat map of point density averaged over all phase spaces for all mice (n=15). Circles show the 95% boundaries of NREM, REM and wake for each mouse. Histograms from all mice are shown along the relevant axis with automatically determined thresholds (*)*. **D**. *Overlap of scoring using thresholds determined on one mouse and applied to a different mouse (n=15 mice). As in B*.

We first quantified this similarity in the same animals between days and between light and dark cycles. We used the thresholds defined for one animal on a given light-cycle to score test data from the same animal on a subsequent light- or dark-cycle. The scoring was then compared with that obtained using thresholds determined from the test data itself. Thresholds for one animal were calculated as the distance from the SWS peak, both for OB gamma activity and HPC theta/delta ratio, to correct for the shifts in overall amplitude that might be caused for instance by changes in recording site (see methods for details).

We found that the observed consistency was sufficient to perform highly accurate scoring on the next day light cycle (average over recordings: 97 ± 0.5%, Cohen’s K: 0.97, n=15, Fig. 5Bi) and during the dark cycle (average over recordings: 96 ± 0.9%, Cohen’s K: 0.94, n=4, Fig. 5Bii) using independently defined thresholds (see methods).

We next compared the phase space used for sleep scoring between animals. We found that after normalizing distributions to the mean NREM values, both OB and HPC distributions were highly reproducible across mice and the independently determined thresholds had very close values (Fig. 5C). Scoring one animal using the thresholds determined for another as above, we found that scoring was also highly reliable (average over recordings: 90 ± 2.5%, Cohen’s K: 0.85, n=15, Fig. 5D).

Finally, since gamma oscillations in the olfactory bulb have been linked with information processing and novelty (Kay *et al.*, 2009), we exposed 8 mice to a novel environment for 15min, during which the animals actively explored. On average only 2 ± 1.1% of the time was misclassified as sleep. This demonstrates that any changes in gamma activity linked to behaviour remain well within the bounds of the wake state as previously defined.

This demonstrates that brain state related changes in gamma power are quantitatively robust over multiple days, throughout the circadian cycle and during exposure to new environment. Moreover, the phase space thus constructed is highly reproducible between animals. This makes it an excellent parameter to use for automatic methods of scoring and a promising tool for comparing sleep in cohorts of animals.

### A powerful tool to study mismatch between brain state and motor activity

A major issue with current approaches to sleep scoring is that EMG activity conflates absence of movement and sleep which suffers from notable exceptions such as during freezing behaviour. Freezing is a widely-studied behaviour in paradigms such as fear conditioning. It is defined as a complete absence of all movement except for respiration. This absence of movement is associated with a strong drop in EMG power. Although it has been shown that average EMG power is lower during sleep than freezing (Steenland & Zhuo, 2009), we investigated whether freezing could be misclassified as sleep using EMG power and whether OB gamma power could resolve this issue. Six mice were therefore fear-conditioned by pairing tones with mild footshocks and during test sessions displayed robust freezing to tone presentation (see methods).

The example session shown in Fig. 6A illustrates the strong expected drop in EMG power during freezing periods, sometimes below the sleep/wake threshold independently determined during a previous sleep session. Although the EMG power is indeed on average higher than during the sleep state, freezing time point can be misclassified as sleep (Fig. 6B). In the example in Fig. 6A, 54% of freezing periods were classified as sleep and EMG shows a similar drop in power at freezing and sleep onset (Fig. 6C). In sharp contrast, gamma power remains systematically above the sleep/wake threshold (Fig. 6B). Gamma power triggered on freezing onset shows that the variable is independent of freezing onset (Fig. 6C).

**Figure 6.**
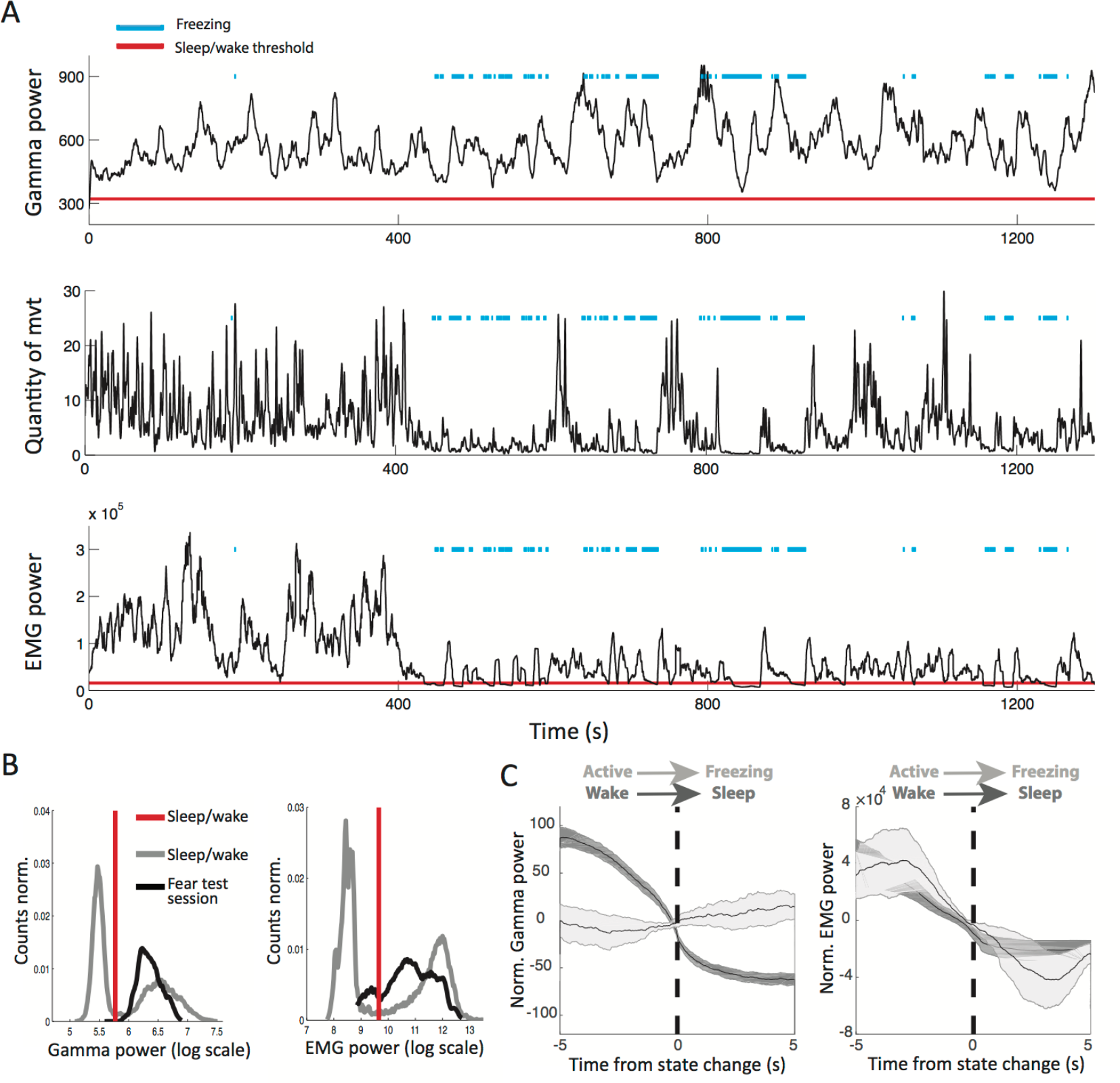
**A**. *OB gamma power, quantity of movement and EMG power during an example test session during which the mouse displayed freezing episodes (blue line). Freezing was determined using the quantity of movement. Red lines indicate the sleep/wake thresholds independently determined during a previous session in the homecage for OB gamma and EMG power*. **B**. *In gray, distribution of gamma (left) and EMG (right) power during the homecage session. In black the distribution during the test session, including the freezing periods. For gamma power all values recorded during the test session are classified as wake whereas the EMG power during values from freezing periods are below the threshold*. **C**. *Averaged OB gamma power (left) and EMG power (right) triggered on two types of transitions from mobility to immobility: the wake to sleep transition and the active to freezing transition. OB gamma power drops when the animal falls asleep but not when the animal freezes. EMG power shows similar changes during freezing and sleep onset (n=6, error bars: s.e.m)*

Freezing is a behaviour that dissociates complete immobility from sleep, allowing us to clearly show that OB gamma power is tracking transitions from wake to sleep and not from mobility to immobility. EMG in contrast is an unreliable marker for sleep scoring when animals are susceptible to display immobility during wakefulness.

### The different dynamics of waking up and falling asleep

Our automated sleep scoring method classifies all data points into periods of wake, NREM and REM. Another essential aspect of sleep study concerns the intermediate transitory period between these two states. Here we show that these transitions can be studied by visualizing moment to moment variations in brain state as a moving point in the previously described phase space (Fig. 5C). The trajectories that this point describes during transition periods can then be studied using classical tools from kinematic analysis.

Shifting to this finer time scale, we replaced the previously used threshold with a transition zone, consistent with many experts arguing that there is no single time point corresponding to the onset or offset of sleep (Ogilvie, 2001). For each point in phase space the probability of remaining in the current state is calculated revealing highly stable zones (Fig. 7A, dark blue) and zones of instability (Fig. 7A, pale colours) in which the state of the brain is changing. Two transition zones naturally emerged along the gamma power axis between sleep and wake states and the theta/delta ratio axis between REM and NREM (grey lines in Fig. 7A for sleep/wake). We propose three measures based on these transitions to study the dynamics of state change and illustrate that they allow for novel characterization of the sleep/wake transition.

**Figure 7.**
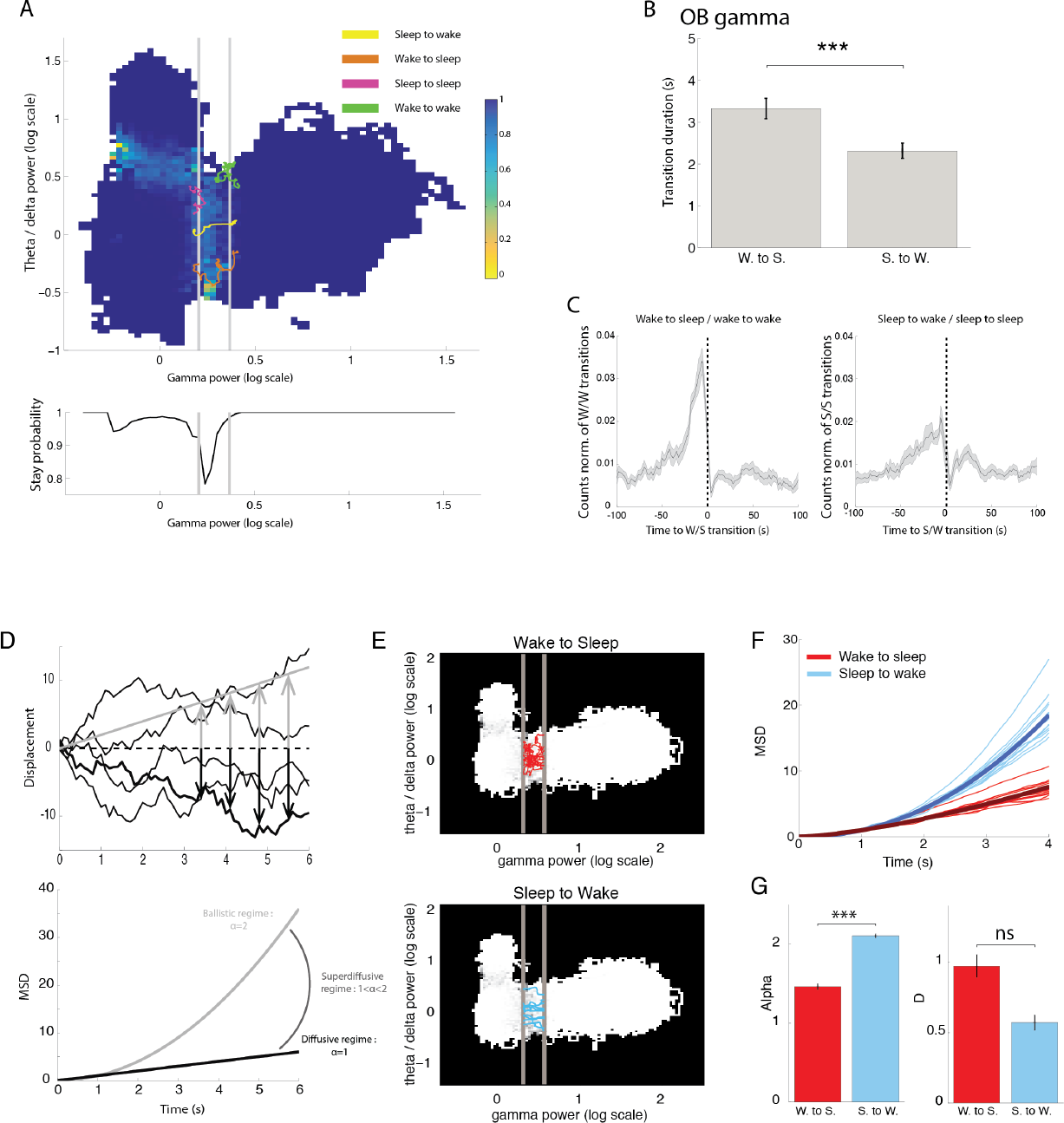
**A** *Phase space in which color indicates the probability of staying in current state in the next 3 seconds. Far from state boundaries, dark blue indicates that the state is unchanging however the frontiers between states are characterized by a lower stay probability. Averaging the stay probability along the gamma power axis shows a strong dip in stay probability that is identified as the transition zone (gray zone). This allows for automatic definition of this zone across mice. The four example trajectories show describe full transitions (sleep to wake and wake to sleep) and aborted transitions (sleep to sleep and wake to wake)*. **B**. *Traversal time of the transition zone defined as in A showing that wake to sleep transitions are significantly longer than sleep to wake transitions (gamma: p=0.0007, n=15, paired t-test). Note that both methods predict approximately the same transition durations. (error bars: s.e.m)* **C**. *On the right cross correlation of aborted transitions times relative to true transitions. Wake to sleep transitions appear more tightly coupled to aborted transitions from wake than sleep to wake transitions with aborted transitions from sleep (higher peak around −30s). (comparison of 20s before and after transition p=0.04, n=15, paired t-test; error bars: s.e.m)* **D**. *Schematic of ballistic and diffusive motion. Ballistic motion linearly relates displacement (position relative to initial condition) to time as shown by the orange line (top). The mean square displacement therefore is related to the square of time (bottom). This type of movement is deterministic: repeated trajectories are all superimposed. On the other hand, diffusive motion is a stochastic process: repeated measures yield varying trajectories (purple lines). The mean square displacement is linearly related to time (bottom). When the exponent of time (alpha) is between 1 and 2 the motion is described as superdiffusive*. **E**. *The phase space of an example mouse is shown with transition zones shown in gray. 10 randomly chose transition trajectories have been plotted from wake to sleep (top) or sleep to wake (bottom)*. **F**. *Mean square displacement for sleep to wake and wake to sleep trajectories as a function of time after normalisation by D for clarity. Thick line: average over all mice (n=15). Note the clear separation of the two sets of lines*. **G**. *The functions in C are fit by the model equation that defines alpha and D. Mean alpha values but not D coefficients are significantly different between the two types of trajectories. (alpha: p=7E-11, D: p=0.2, paired t-test, n=15)*

First, we defined “true transitions” as a full traversal of the transition zone. Interestingly we found that the duration of this traversal depended on the direction. Wake to sleep transitions were significantly slower than sleep to wake transitions (Fig. 7B).

Second, we also defined “aborted transitions” during which the brain state enters this transition zone but does not make a full traversal. Aborted wake/wake transitions were more frequent than sleep/sleep transitions (p=0.005, n=15, paired t-test) and tended to be strongly grouped in the 30s preceding complete wake/sleep transitions (Fig. 7C).

Finally, we adopted a dynamic approach to analyse these trajectories. Differences in transition durations (Fig7. B) from wake to sleep and sleep to wake, given the constant width in phase space of the transition zone, can be interpreted as differences in speed only if the movement is well described by ‘ballistic’ motion. Ballistic motion indicates that time, speed and distance are linearly related (Fig7. D, gray). This is not true for all trajectories, particularly those describing stochastic processes such as Brownian motion (Fig7. D, black). A more general approach proposes describing the movement of the particle using a simple 1D model of its mean square displacement:

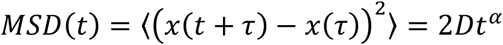

Theory of Brownian motion indicates that when alpha = 2, the movement is ballistic and for alpha=1, the movement is diffusive. For alpha<2 and >1, the movement is described as super-diffusive. Transitions with essentially similar dynamics but different speeds are described by the same value of alpha but different values of D. On the other hand, a change in the value of alpha would argue for different processes underlying the transition.

Fig. 7E compares the same number transition trajectories from wake to sleep (top) and from sleep to wake (bottom) from an example mouse using OB gamma power. The wake to sleep trajectories appear much more convoluted and tile the plane more densely than the sleep to wake trajectories, suggesting that their movement is more diffusive. The MSD for both transition types is shown in Fig. 7F for all mice. The clear segregation of the two sets of curves therefore indicates a difference in alpha.

The MSD of trajectories were well fit using the phenomenological model described above (mean R2 = 0.996, all R2 larger than 0.988). Fitting reveals that the movement from sleep to wake is ballistic (mean alpha = 2.1) whereas the motion from wake to sleep is more diffusive (mean alpha = 1.47) (Fig. 7G). Importantly we observed no significant difference in the diffusion constant D indicating that the difference in transition duration is due to a change in type of dynamics and not a change in this proxy for speed (Fig. 7G).

Overall, using the phase space approach we can clearly demonstrate a strong and novel dichotomy between the sleep-wake and wake-sleep transitions. In particular, the awakening transition is fast and ballistic whereas the process of falling asleep is slower and more stochastic, preceded by multiple failed attempts to transition. Finally, the frequent aborted sleep/sleep transitions that tend not necessarily to precede actual awakenings and may be linked to subthreshold arousal phenomena such as the cyclic alternating pattern [Halasz, 1998].

## Discussion

Sleep scoring is the essential first step in any study of brain states and the characterization of microstructure an essential aspect of understanding sleep’s dynamics in health and disease (Halasz, 1998). Here we establish for the first time a methodology for addressing both issues with brain signals alone. We found that gamma oscillations in the olfactory bulb was a brain signal usable to continuously identify the different sleep stages. It can be substituted to the muscular activity or body movements that are required in all the other sleep scoring methods. It can also be used to analyse the fine dynamics of state transitions by applying phase space trajectory analysis.

The novel sleep scoring method we propose relies on activity recorded in the HPC and the OB only. Implantation of electrodes for recording LFP in these two areas is easy to achieve because both areas show robust oscillations in the theta and gamma ranges respectively at multiple recording sites. After implantation the method is full automated and therefore removes the time-consuming steps of scoring by hand the full data set or a training set to calibrate semi-automatic algorithms. We have shown that this method for sleep scoring is robust to slight changes in implantation site, across days and between different animals. It therefore allows easy comparison between mice and throughout time, and may be between data sets from different laboratories.

Beyond the technical ease of use, this method also provides a promising framework for the study of the dynamics of brain states. Using activity recorded in the brain and not muscle activity allows to track sleep/wake activity independently of movement. This could provide the heretofore lacking methodology to study phenomena such as REM without atonia induced in lesion studies (Lu *et al.*, 2006). Our construction of a highly-reproducible phase space across animals and days allows for easy pooling of data sets and comparison of dynamics. Finally, gamma activity in the OB is a variable with fast dynamics that allows to study fine time-scale transitions not accessible to other, slower sleep-related oscillations such as delta power.

Finally, the mechanisms underlying the sleep switch have been extensively studied (Saper et al., 2010) but how this related to changes in more global physiological parameters is an open but essential question. The process of falling asleep has been described using behavioural, physiological and electrophysiological variables as a progressive sleep onset period in humans (Ogilvie, 2001). After awakening, the prolonged deficits in performing certain tasks named sleep inertia (Marzano et al., 2011) has been described in humans and in rats and mice OFF periods were observed up until 5 minutes after awakening (Vyazovskiy et al., 2014). All these studies demonstrate that transitions should not be considered as points in time but instead as extended periods of time justifying our use of transitions zones.

We advocate constructing phase spaces with electrophysiological variables capable of tracking high speed changes in the brain stat and that are reproducible between animals as an important step in characterizing these dynamics. We have proposed a simple methodological tool kit for capturing the essential aspects of state transitions and shown its efficiency in describing the deep asymmetry between falling asleep and waking up (Magnin et al., 2010).

We suggest that constructing phase spaces with electrophysiological variables capable of tracking high speed changes in the brain stat and that are reproducible between animals constitutes an important step in characterizing these dynamics. We have proposed a simple methodological tool kit for capturing the essential aspects of state transitions and shown its efficiency in describing the deep asymmetry between falling asleep and waking up (Magnin et al., 2010). The description of falling asleep as a slower, diffusive event whereas waking up constitutes a sharper transition is novel. It is however supported by close inspection of the activity changes at transition times in brainstem and hypothalamic nuclei responsible for the sleep/wake switch that show faster dynamics upon awakening than falling asleep (Cox et al., 2016; Takahashi et al., 2006, 2009).

Altogether, these observations raise the question as to why, compared with other areas, the OB is so well suited to identifying the switch between sleep and wakefulness. One possible explanation is the massive inputs from most of the neuromodulator systems and notably those involved in the control of sleep: cholinergic (D’Souza & Vijayaraghavan, 2014, Senut *et al.*, 1989), hypocretinergic/orexinergic (Gascuel *et al.*, 2012) and noradrenergic (Shipley *et al.*, 1985). Moreover, receptors of these neuromodulatory systems are strongly expressed in the OB (McCune *et al.*, 1993; Hardy *et al.*, 2005). In turn, the different neuromodulators have been shown to modulate gamma oscillations (Hall & Delaney, 2001; Gire & Schoppa, 2008; Li & Cleland, 2013).

A second avenue of explanation is functional. Interestingly, olfaction is the only sensory system in the mammalian brain that does not pass through the thalamic relay before reaching the cortex. The thalamus is thought to play an important role in information gating (McCormick & Bal, 1994) during sleep. Given the link between OB gamma oscillation, task demands (Beshel *et al.*, 2007; Martin & Ravel, 2014) and sensory discrimination (Lepousez & Lledo, 2013), their suppression during sleep could provide a mechanism for sensory gating.

A recent method has been proposed to track continuously vigilance states by tracking pupil diameter (McGinley *et al.*, 2015). Interestingly it was showed that pupil fluctuation follows the activity of cholinergic and adrenergic activity in the cortex (Reimer *et al.*, 2016). However, this method is difficult to implement in freely moving animals. The results shown here suggest that the olfactory bulb oscillations could offer an attractive strategy for the monitoring of vigilance states in natural situations.

## Materials and methods

### Subjects and surgery

A total of 15 C57Bl6 male mice (*Mus musculus*), 3–6 months old, were implanted with electrodes (tungsten wires) in the right olfactory bulb (AP +4, ML +0.5, DV −1.5) and in the right CA1 hippocampal layer (AP −2.2, ML +2.0, DV −1.0). 6 of these mice were also implanted with a hooked EMG wire in the right nuchal muscle. 6 mice were also implanted in the right prefrontal cortex (AP +2.1, ML +0.5, DV −0.5) and parietal cortex (AP −1.7, ML +1.0, DV −0.8). One mouse was recorded with a sixteen-site linear probe (100um spacing, Neuronexus Tech, Ann Arbor, MI, USA). During recovery from surgery (minimum 3 d) and during all experiments, mice received food and water ad libitum. Mice were housed in an animal facility (08:00–20:00 light), one per cage after surgery. All behavioural experiments were performed in accordance with the official European guidelines for the care and use of laboratory animals (86/609/EEC) and in accordance with the Policies of the French Committee of Ethics (Decrees n° 87–848 and n° 2001–464). Animal housing facility of the laboratory where experiments were made is fully accredited by the French Direction of Veterinary Services (B-75-05-24, 18 May 2010). Animal surgeries and experimentations were authorized by the French Direction of Veterinary Services for K.B. (14-43).

Signals from all electrodes were recorded using an Intan Technologies amplifier chip (RHD2216, sampling rate 20 KHz). Local field potentials were sampled and stored at 1,250 Hz. Analyses were performed with custom made Matlab programs, based on generic code that can be downloaded at http://www.battaglia.nl/computing/ and http://fmatoolbox.sourceforge.net/.

### Fear conditioning

Habituation and fear conditioning took place in context A consisting of a square transparent Plexiglas box in a black environment with a shock grid floor and cleaned with ethanol (70%) before and after each session. Extinction learning and test sessions were performed in context B consisting of cylindrical transparent Plexiglas walls with a grey plastic floor placed in a white environment and cleaned with acetic acid (1%) before and after each session.

To score freezing behaviour animals were tracked using a home-made automatic tracking system that calculated the instantaneous position of the animal and the quantity of movement defined as the pixel-wise difference between two consecutive frames. The animals were considered to be freezing if the quantity of movement was below a manually-set threshold for at least 2 s.

On day 1, mice were submitted to a habituation session in context A, in which they received four presentations of the CS- and of the CS+ (total CS duration, 30 s; consisting of 50-ms pips at 0.9 Hz repeated 27 times, 2 ms rise and fall; pip frequency, 7.5 kHz or white-noise, 80 dB sound pressure level). Discriminative fear conditioning was performed on the same day by pairing the CS+ with a US (1-s foot-shock, 0.6 mA, 8 CS+ US pairings; inter-trial intervals, 20–180 s). The onset of the US coincided with the offset of the CS+. The CS- was presented after each CS+ US association but was never reinforced (5 CS-presentations; inter-trial intervals, 20–180 s). On day 2 and day 3, conditioned mice were submitted to a test session in context B during which they received 4 and 12 presentations of the CS- and CS+, respectively.

### Histological analysis

After completion of the experiments, mice were deeply anesthetized with ketamine/xylazine solution (10% /1%). With the electrodes left *in situ*, the animals were perfused transcardially with saline (~50 ml), followed by ~50 ml of PFA (4 g/100 mL). Brains were extracted and placed in PFA for postfixation for 24 h, transferred to PBS for at least 48 h, and then cut into 50-μm-thick sections using a freezing microtome and mounted and stained with hard set vectashield mounting medium with DAPI (Vectorlabs).

### Bimodality quantification

Bimodality was quantified by fitting a mixture of two normal distributions and evaluating either Ashman’s D (Ashman *et al.*, 1994)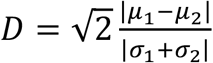 where D>2 is required for a clean separation or the overlap of the two distributions.

### Automatic sleep scoring algorithm

LFP recordings from the OB were filtered in the gamma (50-70Hz) band and instantaneous amplitude derived from the Hilbert Transform. This time-series was then smoothed using a 3s sliding (Fig.1D) window and the distribution of values could be fit with a mixture of two normal distributions. To maximize the probability of correct classification, the threshold between sleep and wake should be defined as the intersection of these two distributions. This value however depends on the amplitude of the two distributions and therefore on the ratio of sleep and wake recorded. To establish a threshold independent of this ratio, the two distributions are normalized to each have area one (Fig2.Bi, right) and the intersection of these distributions is used. Values inferior to this value are classified as sleep and those superior as wake. Periods of sleep and wake shorter than 3s were merged into the surrounding periods to avoid artificially short epochs. Then, LFP recordings from the HPC restricted to the sleep periods defined above, were filtered in the theta (5-10Hz) and delta (2-5Hz) bands and instantaneous amplitude derived from the Hilbert Transform. The ratio of the theta and delta powers was smoothed using a 2s sliding window and the distribution of values was fit by a single normal distribution that accounted for the NREM data points (low theta/delta ratio). The REM/NREM threshold was placed at the point above which the residuals systematically explained more than 50% of the actual data (Fig2.Bii). Periods of NREM and REM shorter than 3s were merged into the surrounding periods to avoid artificially short epochs.

### Automatic EMG scoring

Automatic EMG scoring was performed in a similar fashion to automatic OB gamma power scoring. EMG data was filtered in the 50-300Hz band and instantaneous amplitude derived from the Hilbert Transform. This time-series was then smoothed using a 2s sliding window and the distribution of values could be fit with a mixture of two normal distributions. The intersection of these two distributions, once normalized provided the sleep-wake threshold. The theta/delta power ratio and period dropping procedures are the same as above.

### Manual sleep scoring

Automatic scoring was performed independently by two experimenters using a home-made matlab GUI. The scorers were provided with EMG (raw, filtered in the 50-300Hz band and smoothed instantaneous amplitude) and HPC (raw, low frequency spectrogram and smoothed instantaneous theta to delta ratio) and 3s windows were determined to be NREM, REM or Wake depending on which brain state was judged to be in the majority.

### Evaluation of overlap of scoring methods

The percentage agreement between methods is calculated for each state and shown in the relevant figures. Given that average agreement can be potentially misleading, we also used the confusion matrix to calculate Cohen’s κ (Cohen, 1960) defined as:

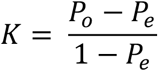

With

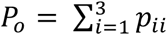 where *p_ii_* is the probability that both methods classify data as the identical state i (REM, NREM, wake).

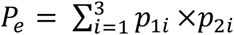 where *p_1i_* and *p_2i_* are the independent probabilities that methods 1 and 2 will classify data as state i.

We applied the same criteria as used in (Libourel *et al.*, 2015) to evaluate the quality of the agreement:

| Quality of agreement | Cohen's $\kappa$ |
| --- | --- |
| Almost perfect | >0.81 |
| Substantial | 0.8-0.61 |
| Moderate | 0.6-0.41 |
| Fair | 0.4-0.21 |
| Slight | 0.2-0 |
| Poor | <0 |

## Acknowledgements

S.B. did the experiments and analysed the data. M.M.L., G.d.L. and J.M.L. did several experiments included in the dataset. H.G. performed the histology. S.B. and K.B. wrote the manuscript with the help of M.M.L and J.M.L. This work was supported by the Fondation pour la Recherche sur le Cerveau (FRC), by the French National Agency for Research ANR-12-BSV4-0013-02 (AstroSleep), by the CNRS: ATIP-Avenir (2014) and by the city of Paris (Grant Emergence 2014). This work also received support under the program Investissements d’Avenir launched by the French Government and implemented by the ANR, with the references: ANR-10-LABX-54 MEMO LIFE and ANR-11-IDEX-0001-02 PSL* Research University. G.d.L. and M.M.L. were funded by the Ministère de l’Enseignement Supérieur et de la Recherche, France. S.B. was funded by the ENS-Ulm, PSL Research University and the Ministère de l’Enseignement Supérieur et de la Recherche, France.

---

**Figure S1.**
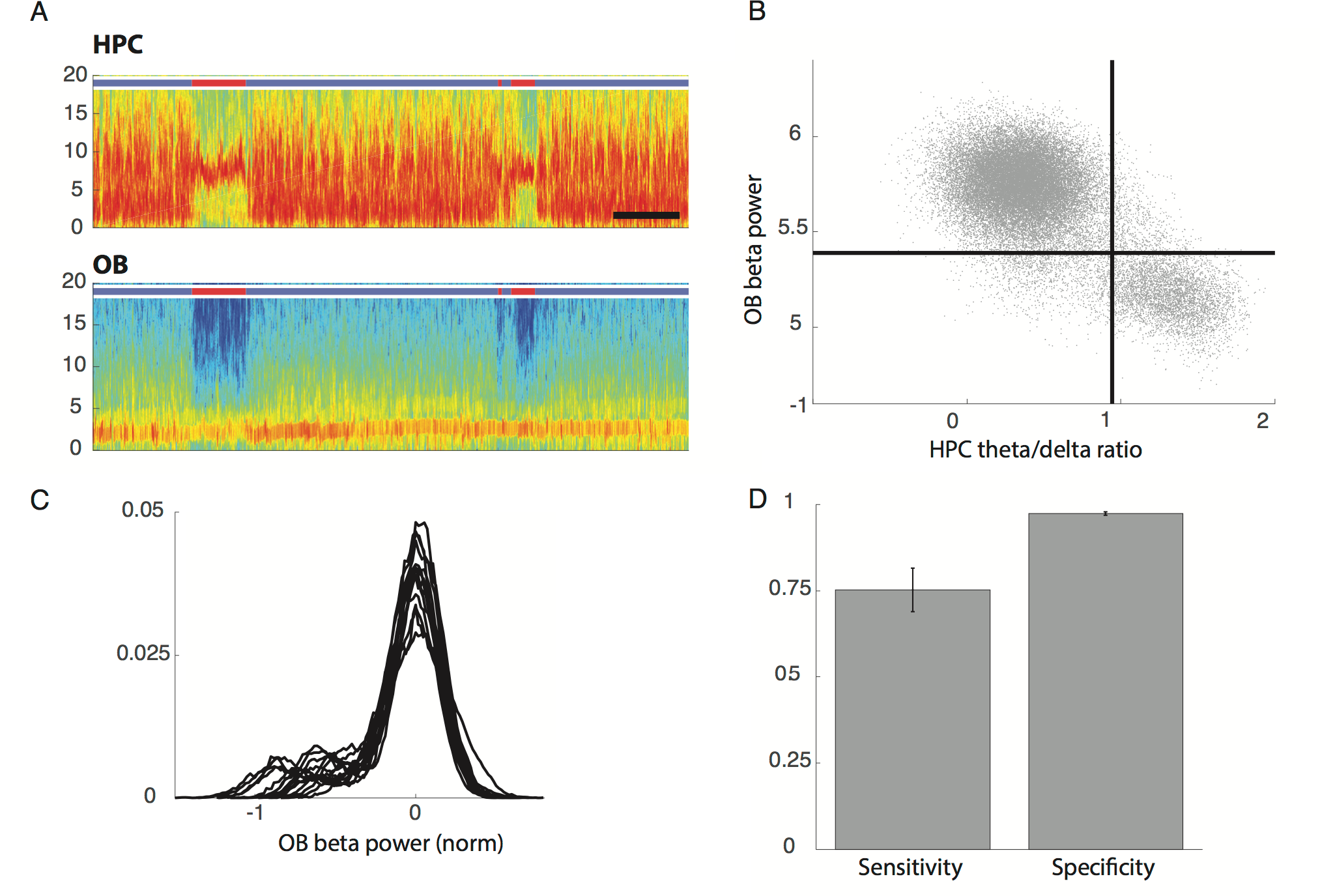
**A**.Example spectrograms of OB (top) and HPC (bottom) activity. The coloured bar indicates the state of the animal defined using OB and HPC activity: blue for SWS, red from REM and gray for wake. Note that during REM periods, defined by strong theta activity in the HPC, there is a drop in a wide band from 10 to 20Hz in the OB. (bar: 3min) **B**.Correlation of OB beta power and HPC theta/delta ratio power for the same mouse as in A with automatically determined thresholds for each distribution in black. Dots in the upper left and lower right hand corner are identically classified by both approaches. **C**.Distributions of beta power in the OB during sleep (n=15). **D**.Sensitivity and specificity of REM identification using OB beta power, where true REM is defined using the HPC activity. Sensitivity is defined as the proportion of ‘true’ or HPCal REM correctly identified as REM and specificity as the proportion of ‘true’ or HPCal SWS correctly identified as SWS. (n=15, error bars: s.e.m)

